# From resting state to inactivation: the dynamic journey of the inactivation peptide in Shaker Kv channels

**DOI:** 10.1101/2024.07.10.602816

**Authors:** Roshan Pandey, Tanja Kalstrup, Rikard Blunck

## Abstract

Shaker Kv channels inactivate rapidly to culminate the action potential and maintain the homeostasis of excitable cells. The so-called N-type inactivation is caused by the first 46 amino acids of the N-terminus of the channel, known as the inactivation peptide. Numerous mutational studies have characterized N-type inactivation functionally, however, the position of the inactivation peptide in the resting state and its transition during inactivation is still debated. Here, we tracked the movement of the inactivation peptide during inactivation using voltage clamp fluorometry. By inserting an unnatural amino acid, 3-[(6-acetyl-2-naphthalenyl) amino]-L-alanine (Anap), which is sensitive to changes in environment, we identified the movements of ball and chain separately. Our data suggests that N-type inactivation occurs in a biphasic movement by first releasing the IP, which then blocks the pore from the cytoplasmic side. To further narrow down the resting position of the inactivation peptide, we used Lanthanide-based Resonance Energy transfer and transition metal (tm)FRET. We propose that the inactivation peptide is located in the window formed by the channel and the T1 domain, interacting with the acidic residues of the T1 domain.

## Introduction

During an action potential, voltage-gated potassium (Kv) channels open in response to membrane depolarization and allow the passage of K^+^ ions to repolarize the membrane. Duration and pattern of action potentials in excitable cells is partially regulated by fast inactivation during which the cytosolic N-terminus of Kv channels binds to the open pore and blocks the entry of ions (N-type inactivation) (1-5). Fast inactivation is observed in Kv 1.4, Kv 3.4, Kv 4.2 and is caused by a peptide at the N-terminus (inactivation peptide, IP). Variably, coexpression with β-subunit imparts inactivation to the non-inactivating currents of Kv1.1, 1.2, 1.3 and 1.5 channels (6, 7). Fast inactivation in voltage-gated sodium channels was originally also thought to undergo a similar mechanism, with the IFM motif in the interdomain linker acting as the inactivation peptide. However, high-resolution structures of several Nav channels confirmed that the motif is binding laterally to the pore and an alternative mechanism has been proposed (8-12).

The IPs of N-type inactivating channels have no sequence similarity, but they all contain a hydrophobic tip region of approximately the first 10 amino acids, which in the Shaker Kv channel, have been suggested to inactivate the channel by interacting with hydrophobic pore-lining residues (1, 5). The downstream hydrophilic region which carries a net positive charge, is responsible for increasing the diffusion rate of the IP towards the pore region via long-range electrostatic interactions (5, 13-15).

Below the transmembrane region of Shaker-related Kv1 channels one finds the cytoplasmic T1 domain, which is responsible for subunit tetramerization (16-18). The T1 domain is attached to the first transmembrane helix S1 via the T1-S1 linker in such a way that four windows form between the T1 domain and the transmembrane domain (Fig. 1A). It has been suggested that the N-terminus enters the window of its own subunit in order to reach the inner pore cavity (19, 20). The T1-S1 linker contains conserved negative charges important for accommodating the positively charged chain region (13, 19, 21).

**Figure 1.**
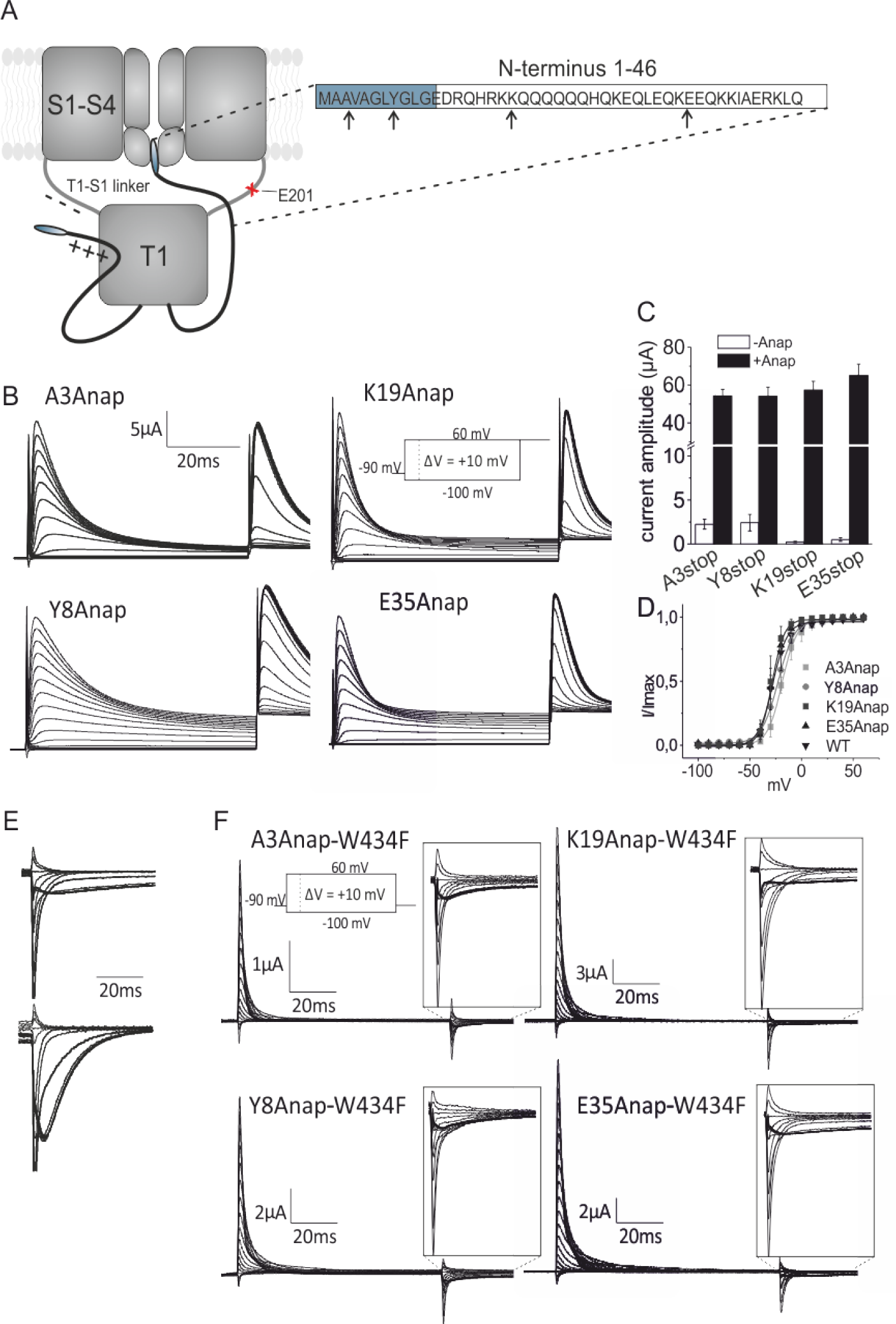
Functional expression with Anap in N-terminus. **A)** Cartoon of the Shaker channel inactivated by the N-terminus. Only two subunits are shown for clarity. Residues used in this study are marked arrows in the N-terminus and a red cross in the T1-S1 lin. **B)** Currents obtained from each N-terminal mutant upon a standard activation step protocol followed by a second pulse to +60 mV. **C)** Expression levels shown as current peak amplitudes upon depolarizing from -90 mV to +40 mV for each mutant in the absence and presence of Anap. Adapted from Kalstrup and Blunck 2015 [17] **D)** IV curve for inactivation obtained by plotting the peak current during the second pulse in at +60 mV in C). **E)** Off-gating currents obtained with W434F in WT full length (upper) and Δ6-46 channels (lower). **F)** Gating currents obtained during a standard activation step protocol. Insets show the presence of charge immobilization.

Even though the mechanism of N-type inactivation is functionally well characterized, any dynamic information about the pathway by which the IP travels from resting state to inactivation is lacking. By inserting a fluorescent unnatural amino acid (Anap) into the IP, using the amber stop codon suppression technique (22-24), we were able to probe relative movements using voltage clamp fluorometry (VCF). This procedure allowed us to compare tip motion (A3 and Y8) and chain motion (K19 and E35). Anap was also inserted into the T1-S1 linker receptor site (E201) to probe the arrival of the IP through the T1 window.

The current model for N-type inactivation generally implies that the N-terminus diffuses in the cytosol and enters through the side windows upon depolarization. Our results, in contrast, suggest that the N-terminus is already bound near or at the T1 windows. In order to probe the resting state position of the IP, we performed lanthanide resonance energy transfer (LRET) experiments by labelling the alpha subunit of the channel (top of S4 and S5 helix) with a donor and the ball (Y8) and chain (E35) peptides with an acceptor fluorophore. To further narrow down the position, we performed transition metal FRET (tmFRET) taking advantage of its sensitivity for shorter distances (25). To this end, we inserted dishistidine motifs at several positions around the T1-S1 window for binding cobalt. By calculating the energy transfer between cobalt and Anap incorporated at positions Y8 and E35 in the IP, we were able to locate the position of IP in the resting state.

## Results

### Anap incorporation does not inhibit N-type inactivation

In order to probe movements of the IP during N-type inactivation, Anap was inserted into selected sites in the Shaker channel via the amber codon suppression technique using the Anap specific tRNA synthetase/tRNA (RS/tRNA) pair (22, 23). To this end, two positions in the tip region, A3 and Y8, and two positions in the chain region, K19 and E35, were used as sites for Anap incorporation (Fig. 1A). Since N-terminal insertion of stop codons can lead to translational reinitiation at downstream start codons, nine start codons were silently mutated as previously described (26). Anap was successfully incorporated into the four positions as determined from the robust expression in presence of Anap and poor expression in the absence of Anap (Fig. 1B-C). All mutants exhibited functional N-type inactivation (Fig. 1B) with WT-like inactivation voltage dependencies (Fig. 1D).

It has previously been shown that when the IP binds to the pore in the inactivated state, the voltage sensors are unable to return to their resting state, making unbinding of the IP the rate limiting step upon repolarization (27). In W434F channels, where ionic currents are blocked but the gating machinery remains normal (28), voltage sensor immobilization materializes as a slow off-gating current during repolarization which disappears in the Δ6-46 deletion mutant (Fig. 1E) (27). We used this feature to verify whether the tip properly reaches the final docking site in the Anap mutants. Indeed, charge immobilization was unaffected in all mutants as shown by the presence of a slow component in the off-gating currents with a time course similar to the one found in WT (Fig. 1F). The finding that the IP reaches the final docking site agrees with the functional current phenotypes seen during N-type inactivation (Fig. 1B). These results show that the presence of Anap in the tip and the chain region did not inhibit the ability of the IP to reach the pore and block the ionic currents.

### Anap in the tip region probes the final step in N-type inactivation

A3Anap and Y8Anap exhibited small voltage dependent fluorescence changes at expression levels above 50 μA when depolarizing to 40 mV (Supplementary Fig S1A-B; the slow inactivation kinetics seen in figure S1A-B are caused by channel clustering) (29). The presence of two fluorescence components of opposite direction made the analysis of A3Anap and Y8Anap challenging due to low dF/F values. In order to enhance the fluorescence signal, we therefore decided to use the non-conducting Shaker-W434F mutant (30), which allows higher expression levels than the conducting channels. The fluorescence changes in W434F channels were similar to those of the conducting channels and did also exhibit two components of opposite direction (Fig. S1C-D and Fig. 2A-B, upper traces). In order to verify whether the fluorescence changes were caused by binding of the tip region to the pore, we used the Kv channel blocker 4-aminopyridine (4-AP), which inhibits the final transitions during channel activation (31-33). Since pore opening needs to occur for the IP to reach the final docking site, we were able to identify fluorescence changes sensitive to the ability of the IP in blocking the pore. Indeed, application of 5 mM 4-AP resulted in inhibition of the second fluorescence component, suggesting that the final blocking step is monitored by Anap (Fig. 2A-B, middle traces).

**Figure 2.**
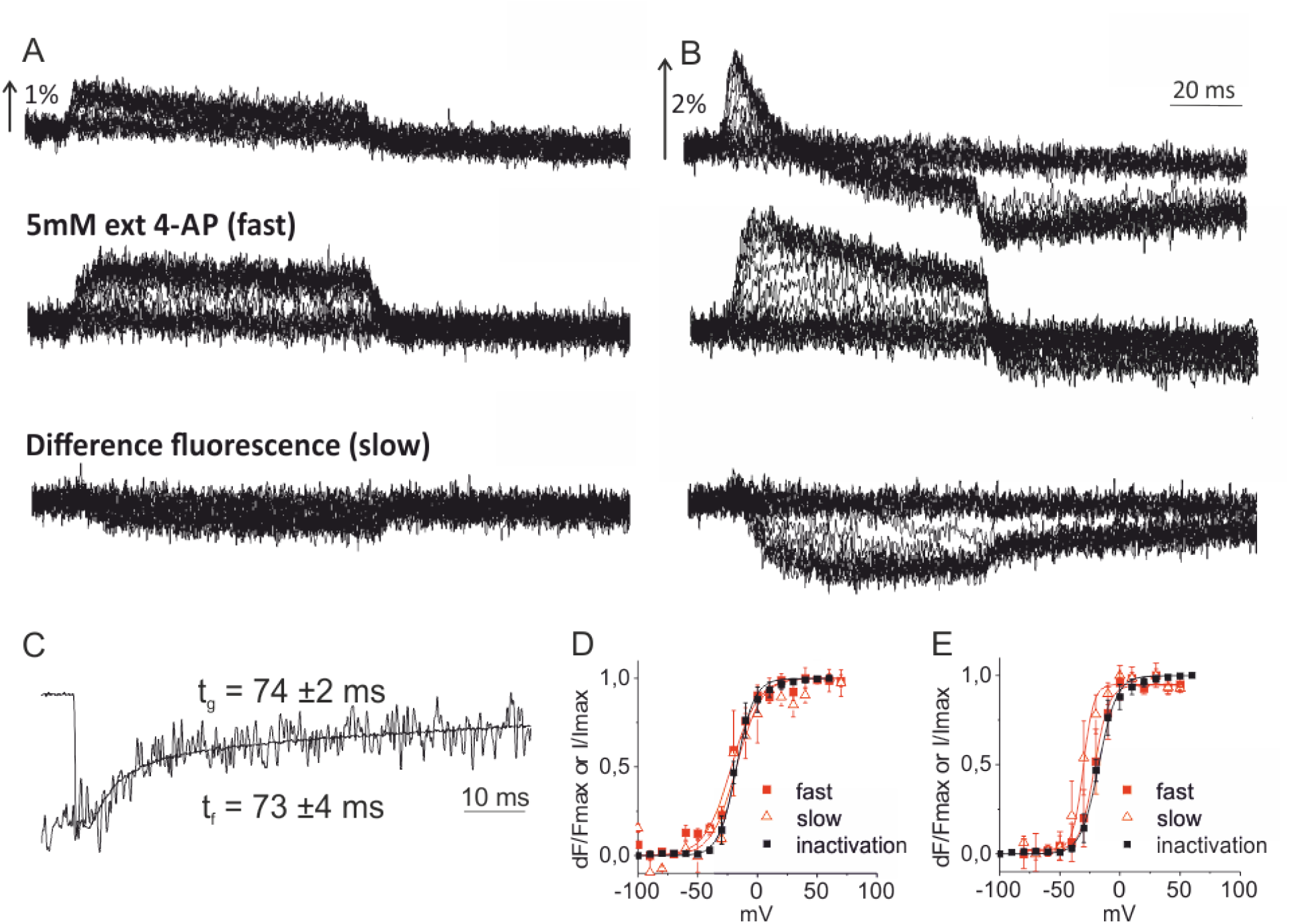
VCF results for the tip region mutants A3Anap and Y8Anapv. **A-B)** Voltage-dependent fluorescence changes obtained during recordings of the gating currents inFig. 1F for A3Anap-W434F A) and Y8Anap-W434F B). Middle traces are obtained in the presence of 5 mM external 4-AP, and the lower traces are the 4-AP sensitive components obtained via subtraction **C)** Overlap of Y8Anap-W434F off-gating current and fluorescence time course during repolarization from 0 mV to -90 mV. **D-E)** A3Anap-W434F (D) and Y8Anap-W434F (E) FV curves of the 4-AP insensitive (slow) and insensitive (34) components plotted together with the IV of inactivation from Fig. 1D.

We obtained the slow 4-AP sensitive component by subtracting the signal in presence of 4-AP from the original signal (Fig. 2A-B, lower traces). In Y8Anap, the fluorescence time course upon repolarization followed the slow kinetics of charge immobilization with a time constant τ_f_= 73 ± 4 ms and τ_g_ = 74 ± 2 ms, respectively, obtained from exponential fits (Fig. 2C). This indicates that the slow fluorescence component directly probes conformational changes during the final blocking step in agreement with its 4-AP sensitivity. The 4-AP sensitive component in A3Anap was too small to confidently fit to an exponential curve and was therefore not analyzed kinetically. Plotting the 4-AP insensitive (34) and 4-AP sensitive (*slow*) component as a function of voltage (FV) for both mutants, revealed that, for both components, the voltage dependence is similar to that of inactivation (Fig. 2D-E). Taken together, these results strongly suggest that Anap at position A3 and Y8 in the IP probes a movement which causes the docking of the tip into the final receptor site.

The fast fluorescence component seen here also shows that IP starts to move with the activation of the channel. So, the IP is able to detect channel activation and mechanism of inactivation starts simultaneously with the opening of the channel. To further trace the movement of IP, we decided to insert Anap in the chain region.

### Anap in the chain region probes movements preceding final block

For the chain mutants K19Anap and E35Anap, we followed the same protocol. In order to optimize the fluorescence changes, we required high expression levels (Fig. 3A-B). In contrast to the tip mutants, they exhibited a single fluorescence component, which made the fluorescence signal easier to analyze and thus excluded the need for using W434F. In contrast to the tip mutants, K19Anap and E35Anap produced fluorescence intensity signals insensitive to application of 4-AP (Fig. 3C-D), suggesting that they report movement that occurs prior to the IP entering the pore. The FV of K19Anap correlated with the IV curve for inactivation with midpoint values of -19.3 mV and -14.3 mV respectively, as obtained from Boltzmann fits (Fig. 3F). On the other hand, the FV of E35Anap was more shifted towards polarized potentials with respect to inactivation with midpoint values of -24.0 mV and -9.3 mV, respectively (Fig. 3H). The kinetics of inactivation and fluorescence varied from oocyte to oocyte with slower inactivation at increased expression as mentioned earlier (an effect which disappears with W434F, Fig 3E). We therefore could not directly compare the mean of the time constants.

**Figure 3.**
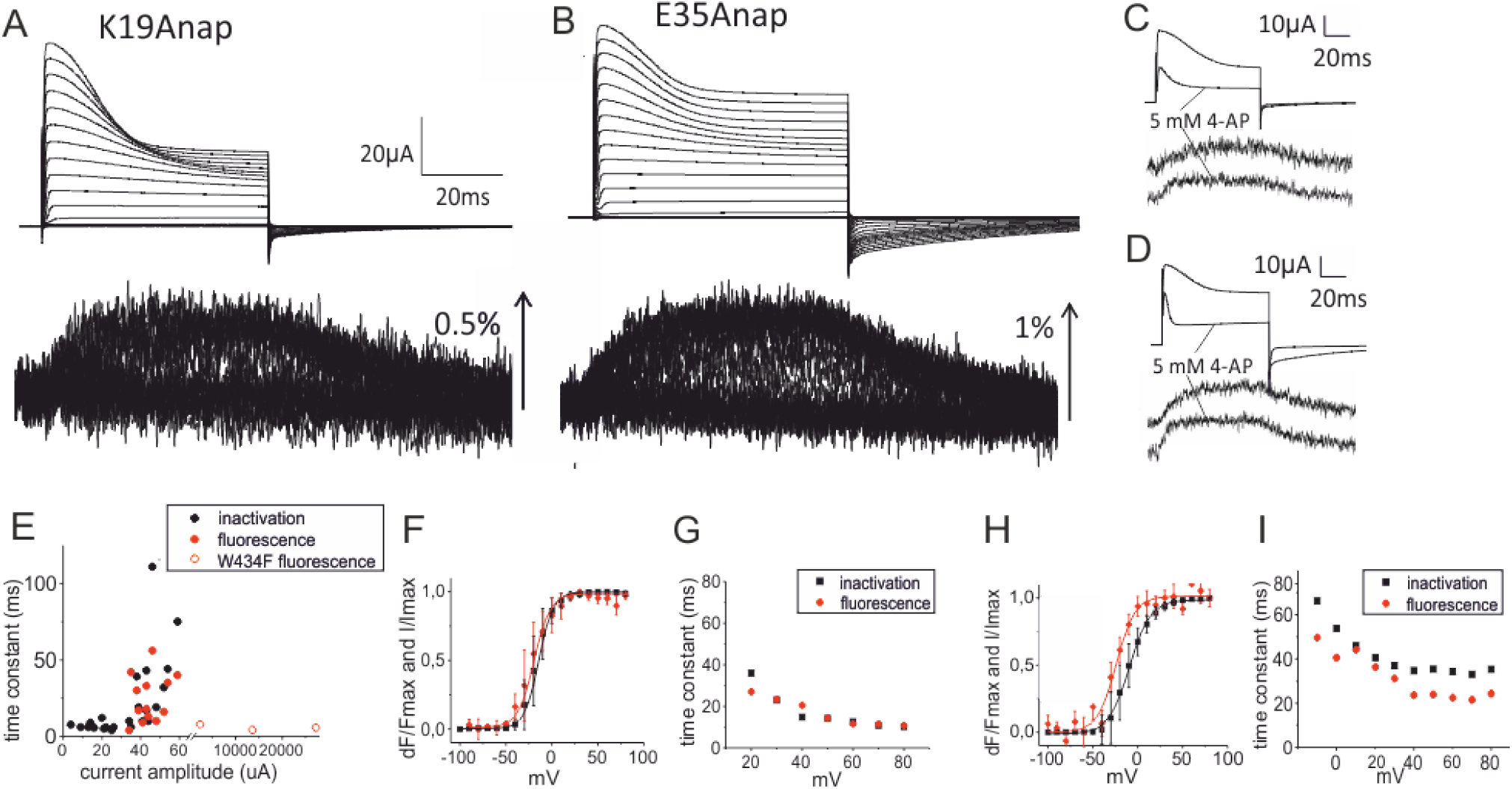
VCF results for the chain region mutants K19Anap and E35Anap. **A)** and E35Anap **B)** and their respective fluorescence changes. **C-D)** Comparison of currents and fluorescence changes before and after application of external 4-AP during depolarization from -90 mV to + 40 mV. **E)** Time constants obtained from exponential fits of E35Anap current inactivation and fluorescence changes during depolarization from -90 mV to +40 mV, are plotted as a function of current amplitude (expression level). The corresponding ionic current of W434F gating currents were calculated using 13 elementary charges per channel [25, 26] with a unitary conductance of 20 pS. **F)** FV (red) and IV (black) curves obtained from K19Anap expressing oocytes. **G)** Time constants obtained from exponential fits of inactivation currents (black) and fluorescence (black) time courses of K19Anap expressing oocytes. H-I) Same as F-G) but for E35Anap expressing oocytes.

Nevertheless, when comparing inactivation and fluorescence kinetics of single oocytes, the data suggests that fluorescence and inactivation exhibit similar kinetics in K19Anap, whereas, in E35Anap, the time constant of fluorescence is approximately 10 ms faster than the one of inactivation (Fig. 3G and I). Taken together, the results indicate that, in both mutants, Anap is probing a transition which precedes final block, due to their 4-AP insensitivity. However, the two movements are not necessarily the same. The K19Anap movement develops closely with inactivation whereas E35Anap fluorescence signal is both shifted to more polarized potentials and faster than inactivation. This indicates a sequential transition mechanism; first, the downstream chain region movement (E35) sets in motion, which then is followed by a second movement involving the upstream part of the chain (K19).

### Location of the inactivation particle during the resting state of the channel

Our fluorescence data suggested a coordinated movement of the ball and chain to cause inactivation of the channel. It senses channel activation and thus has to be in close proximity to the transmembrane domain during the resting state of the channel. This raises the question of where the inactivation particle and the transmembrane region interact during the resting state. To identify this position, we probed whether we could detect the movement of the inactivation peptide in the transmembrane domain both during activation and inactivation. To this end, we inserted Anap at position E201 in the T1-S1 linker (red mark, Fig. 1A). Like the N-terminal mutants, E201Anap exhibited functional N-type inactivation (Fig. 4A) and displayed fluorescence changes at high expression levels (Fig. 4B). As the T1-S1 linker is situated close to the voltage sensor, it is possible that the fluorescence changes of E201Anap originate from conformational rearrangements in the transmembrane domain during channel activation and not from the inactivation peptide. To verify this, the experiments were carried out in presence and absence of the inactivation peptide (inactivation-removed IR, lacking the N-terminus (Δ6-46)). IR-E201Anap channels exhibited no fluorescence changes (Fig. 4C), demonstrating that the movement, probed by Anap in E201Anap is caused by N-type inactivation. In agreement with this, both the V_½_ values and the time constants of the fluorescence signal correlated well with the corresponding values of the decay of the ionic current during fast inactivation (−21.6 mV and -15.8 mV, respectively; Fig. 4D-E). Despite this agreement with the kinetics and voltage dependence of fast inactivation, the fluorescence change was not sensitive to application of 4-AP (Fig. 4F), indicating that after release of the inactivation particle from its resting position during activation, it anneals to the window formed by the T1-S1 linker, the TM and T1 domains before the final step of entering and blocking the pore.

**Figure 4.**
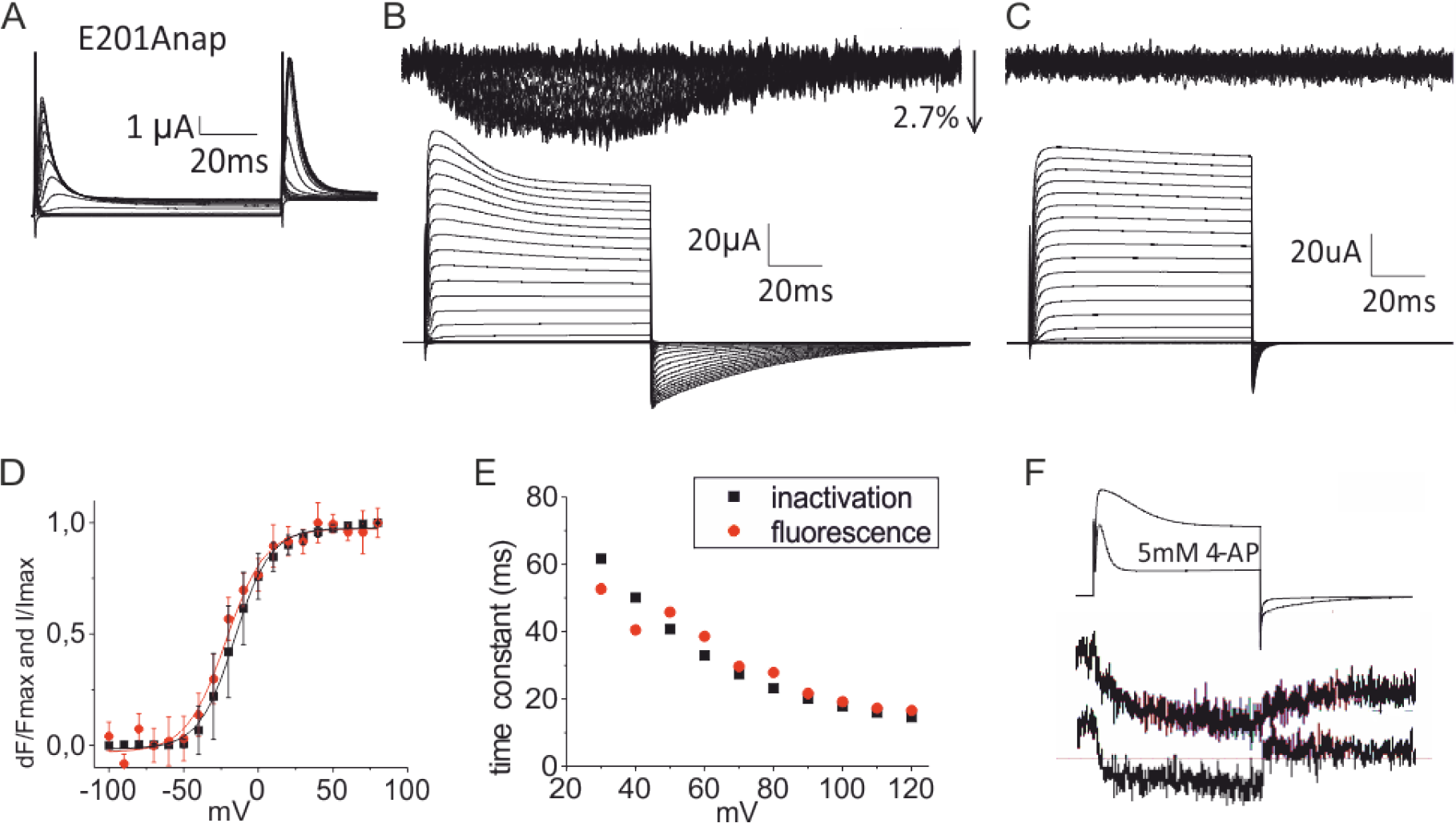
VCF results for the receptor site mutant E201Anap in the T1-S1 linker. **A)** Currents obtained from an E201Anap expressing oocyte from a step protocol as in Fig. 1B. **B)** Example of an oocyte expressing E201Anap showing fluorescence changes upon a standard depolarization protocol. **C)** Same protocol as in B except that the oocyte is expressing an E201Anap mutant in which the N-terminus has been removed (Δ6-46). **D)** Comparison of FV (red) and IV for inactivation (black) of E201Anap. **E)** Time constants of exponential fits obtained from the oocyte shown in B. **F)** Currents and fluorescence during a depolarization step from -90 mV to +40 mV before and after application of external 4-AP.

While the results of E201Anap show that the inactivation particle interacts with this position independent of the entry and block of the pore, the correlation of the kinetics and voltage dependence with inactivation suggests that it is not the resting position of the inactivation particle, as the biphasic signal found for the tip-mutants is missing here. Scanning of the transmembrane region is challenging because many of the positions give strong fluorescence signals during activation independent of fast inactivation. One way to reconstruct the IP movement is to specifically insert tryptophans along putative binding sites (T1 domain, T1-S1 linker) in the background of a tip mutant and a chain mutant (eg. Y8Anap and E35Anap) to induce quenching (or unquenching) of Anap. Since quenching by tryptophan requires contact formation and depends on van-der-Walls radii (35), the method is useful for determination of short range distances (36, 37). Such experiments would allow accurate estimates of the position of the IP with respect to the T1 domain, especially during the resting state which is unknown. Moreover, the direction of the fluorescence change would allow us to determine whether the IP moves towards or away from the inserted tryptophans. To this end, we inserted trp residues in the T1 domain at positions G159W, K178W, E192W, I470W and A391W. However, no quenching of Anap was observed for these positions. We therefore switched strategy: instead of scanning for the resting position, we employed resonance energy transfer to triangulate the exact position.

To probe the position of IP in the resting state, we performed Lanthanide Resonance Energy Transfer (LRET). LRET is a variant of Förster Resonance Energy Transfer (FRET) that uses as donor a lanthanide, in this case terbium. Since terbium does not have a continuous dipole moment, the emission is fully isotropic. This and its long luminescence lifetime (∼1ms) allow a more accurate distance determination compared to FRET (38). The donor was linked to the channel by labelling at position A359C and F425C with Tb^3+^-chelate via a maleimide linker (Fig. 5A) as described earlier (39, 40). The labelling of acceptor positions (Y8 and E35) was challenging as these positions are present on the cytoplasmic side of the channel and therefore, conventional fluorophore labelling using cysteine residues was not possible without avoiding non-specific binding. Anap, on the other hand, did not overlap sufficiently with the Tb^3+^ emission spectra to obtain any meaningful transfer. To overcome this problem, we incorporated an unnatural amino acid, trans-cyclooct-4-en-L-Lysine (TCO), using the amber stop codon suppression technique (24, 41). TCO offers fast, highly specific labelling of fluorophores carrying tetrazine group through strain-promoted inverse electron demand Diels-Alder cycloaddition (SPIEDAC) (42). For acceptor labelling, we used cy3-tetrazine, which shows good spectral overlap with terbium with a R_0_ of 61.2 Å (38). Oocytes expressing the channels without TCO incorporated were used to correct for background correction (see methods for details).

**Figure 5.**
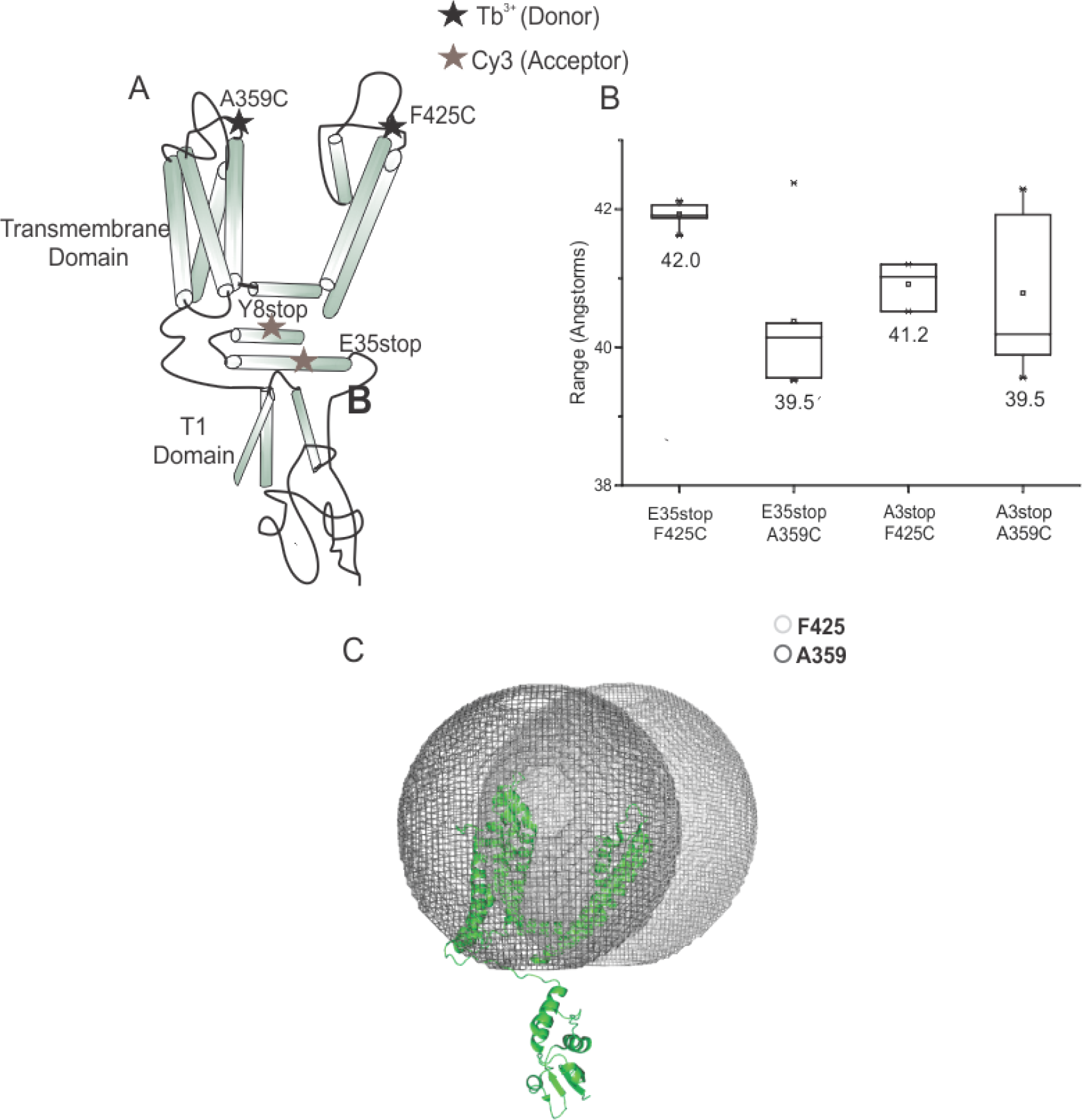
LRET measurement to probe the resting state of the channel. A) Labelling positions for the binding of donor (A359C and F425C) and acceptor fluorophores (Y8Anap and E35Anap). B) Box plot showing distances obtained for different labelled positions in the channel. C) Cartoon representation of the channel monomer with the spheres superimposed on it. The overlapping part of the spheres below the pore represents the probable region for the IP in the resting state.

From the fluorescence decays of donor emission in presence and absence of the acceptor, we calculated the four donor and acceptor distances for the four combinations between two donor and two acceptor positions. By triangulation, we then derived the resting position of the inactivation particle (Fig. 5B): for the donor position A359Tb-Chelate, the resting position was found on a sphere with a radius of 39.5 Å and 40.4 Å for Y8stop and E35stop respectively. Moving the donor to F425Tb-Chelate resulted in a sphere with slightly longer radius of 41.2 Å and 42 Å for Y8stop and E35stop, respectively. The resting position has to be located on the intersecting circle of the two spheres obtained for these positions (Fig. 5C). Most of the circle can be excluded because it is located within the transmembrane region of the channel or the extracellular space. It only remained the region framed by the transmembrane region and the T1 domain close to the pore entry as possible resting state position of the inactivation particle as (Fig. 5C). This result indicates that the IP is prebound to the alpha subunit at that position.

### Transition metal FRET shows distinct movements of ball and chain

To further narrow down the position of the inactivation particle, we required smaller R_0_ such that the energy transfer becomes more sensitive to short distances. While LRET combinations for short distances exist, labeling at the cytosolic surface of the channel with thiol-reactive chemistry is not specific enough. Therefore, we chose transition metal (tm)FRET experiments (43, 44). In tmFRET, a fluorescent dye is used as donor, whereas the terbium ion is replaced with a transition metal ion, but now as an acceptor, with Anap acting as donor. Due to its multivalence, transition metals such as nickel, cadmium or cobalt can be coordinated by dihistidine motifs introduced at the desired position of the protein. Transition metals typically absorb in the visible range, which gives the characteristic blue and pink colors of Ni- and Co-solutions, respectively. The small spectral overlap between Anap emission and cobalt absorption result in a small R_0_ of 12 Å (45-47). Due to the much shorter lifetime of Anap in the nanosecond range, we determined energy transfer efficiency from the fluorescence intensities. (25, 48).

In order to confirm the quenching of Anap fluorescence by cobalt ions, we introduced the histidines in the T1-S1 linker (I197H & K198H) and Anap at position E194. As these positions are very close to each other in the crystal structure of Kv channel (PDB ID 2R9R), we expected high energy transfer between these positions. Indeed, we observed significant energy transfer (dF/F = 15%, Fig 6A, upper trace) and quenching of Anap fluorescence, confirming efficient Anap incorporation at the desired site as well as energy transfer between Anap and cobalt bound to dihistidine motifs.

**Figure 6.**
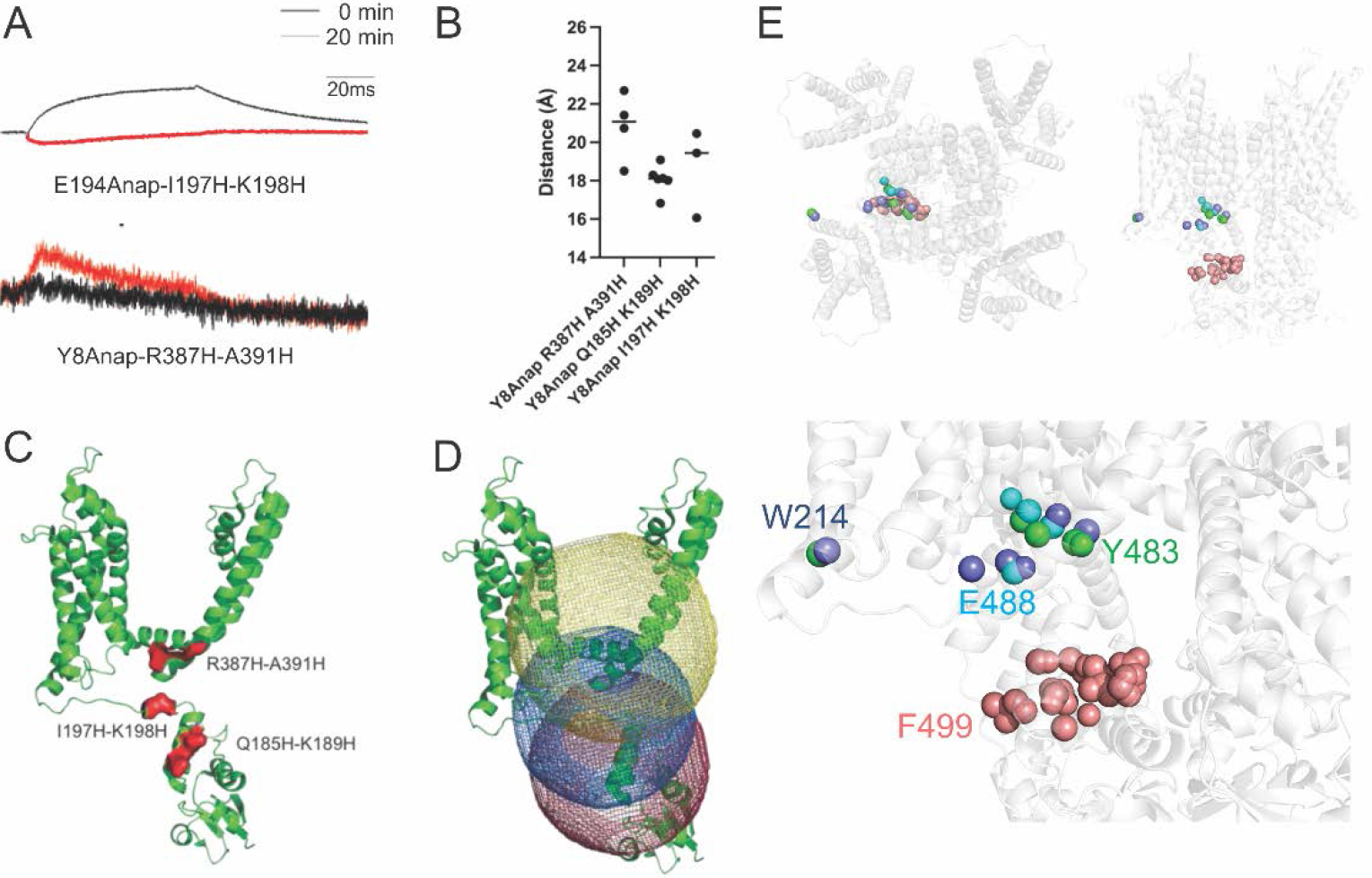
tmFRET results. A) Fluorescence traces recorded for the control E194stop (upper trace) and one of the mutant Y8stop-R387H-A391H (lower trace). The black curve represents the recording at 0 min. Red traces show fluorescence quenching due to energy transfer after the addition of cobalt. B) Calculated distances from tmFRET measurements for the positions mentioned in Fig. 6C. C) dihistidine mutations introduced at various positions in the channel for the binding of cobalt. D) Cartoon representation of the channel monomer with the spheres plotted using the distances shown in 6B (Yellow represents R387H-A391h, blue represents I197H-K198H and red represents Q185H-K189H). The overlapping part of the spheres below the pore represents the region for the IP in the resting state. E) Combined results from LRET and tmFRET: The colored atoms are those that fulfill the distance constraints from the LRET and tmFRET measurements. Y8 should be in contact with one of the red ones, A3 with the cyan or blue ones and E35 with the cyan or green ones. (Model based on PDB: 7SIP (49) with missing regions complemented with alphafold3).

We next introduced cobalt labeling sites near the region identified in the LRET measurements in the window formed by transmembrane region and T1 domain. We introduced di-histidine motifs (1) in the C-terminus of the S4-S5 linker (positions R387H-A391H), (2) in the T1-S1 linker closer to the N-terminus (I197H-K198H) and (3) in the upper half of the T1 domain (positions Q185H-K189H; Fig 6C). Anap was introduced at position Y8 in the ball region of the IP. FRET efficiencies were calculated after correcting for unspecific effects of cobalt, which were then used to calculate the distances. The distances calculated from the energy transfer for different positions are shown in Fig 6B.

One of the limiting factors in our experiment was accessibility of histidines to cobalt ions in the solution. We used the distances from the above-mentioned positions to create intersecting spheres as for the LRET results above (Fig. 6D). The window identified by the spheres for the presence of IP coincided the one observed in LRET measurements. This further confirms our finding that IP resides in the window between the channel and the T1 domain.

Combining the results from the LRET and tmFRET measurements, we can identify those atoms in the intersection of the spheres that fulfill all conditions (Fig. 6E). Assuming that the inactivation peptide folds as an alpha-helix, Y8 points towards the T1-domain (red-colored atoms), whereas A3 points towards the C-terminus of the S6. E35 is likely located close to the S1 (W214).

## Discussion

By genetically incorporating a fluorescent unnatural amino acid on the cytoplasmic side of the Shaker Kv channel, we were able to directly probe conformational changes underlying N-type inactivation. The experiments provide the first structural dynamic information about the IP movement during N-type inactivation. The VCF data suggest a sequential-step model during which the IP undergoes at least two transitions, which is in agreement with previous functional studies that suggest an initial binding of the IP to the T1 domain (19), followed by a second binding to the inner pore (5). Based on our results, we suggest that, upon depolarization, both tip and chain region initially move (transition 1 in Fig. 7). Relative to inactivation, the movement of the downstream chain region (E35) preceded the movement of the upstream chain region (K19), suggesting presence of two pre-inactivation steps. A final movement is responsible for docking the IP tip into the pore, and this movement only involves the tip region (A3 and Y8, transition 2 in Fig. 7).

**Figure 7.**
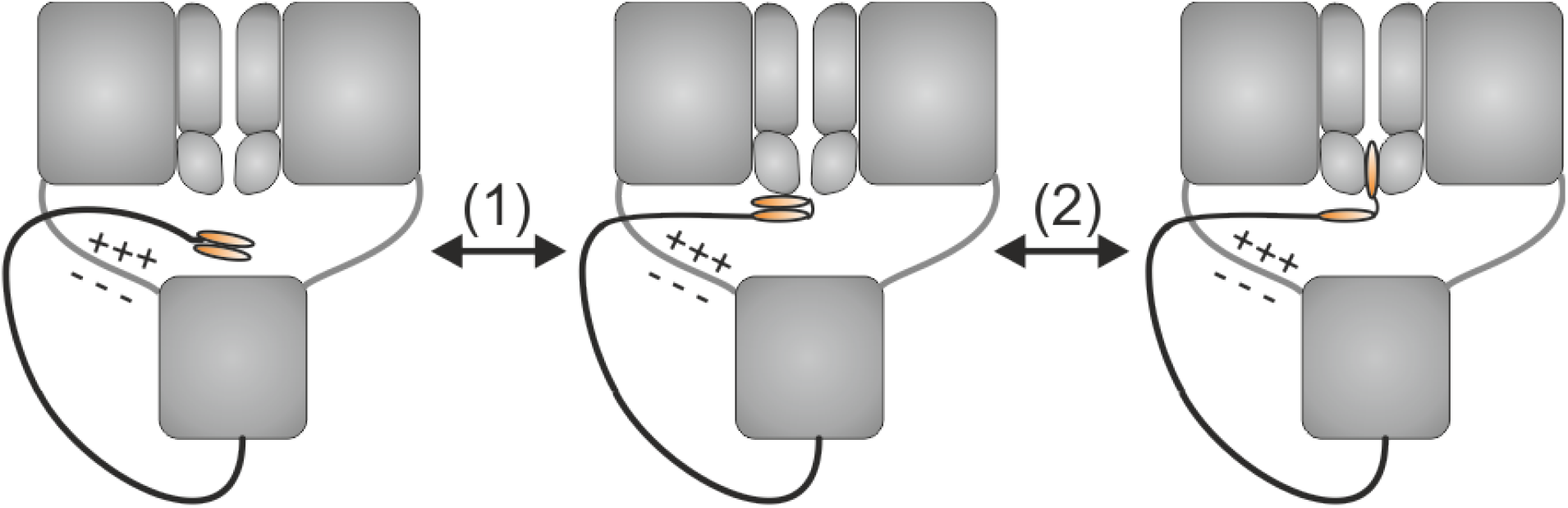
Proposed model for N-type inactivation. In the resting state, IP sits between the channel and the T1 domain interacting with the T1-S1 linker. Binding of the IP to the open pore involves at least two conformational changes. The first involves both tip and chain (1), whereas the second only involves the tip (2).

The finding of two pre-inactivation steps is consistent with previous work on Kv1.4 channels, where a downstream intra-chain electrostatic interaction was identified and suggested to be responsible for initially shortening the distance between the IP and the T1-S1 linker (13, 50). It is therefore possible that position E35 in Shaker lies within a region that is part of a similar interaction as an initial step upon depolarization.

It is worth mentioning that the E201Anap and K19Anap fluorescence data showed similar inactivation profiles. This suggests that Anap experiences the same transition at both positions (i.e., E201Anap probes the reception of K19, and K19Anap probes the movement of the IP towards E201). This interpretation agrees with findings in two other Shaker-related Kv channels where the three conserved glutamates in the T1-S1 linker (199-EEE-201) have been shown to interact electrostatically with positive charges in the early region of the chain which corresponds to the 17-RKK-19 region in Shaker (R18 in AKv1 and 26-RARERER-32 in Kv1.4) (13, 21). To corroborate this further, we performed resonance energy transfer measurements to triangulate the position of the IP during the resting state of the channel. By measuring the distances from the extracellular surface of the channel (top of helix S4 and S6) as well as from the cytosolic surface of the transmembrane region (S4-S5 linker and T1-S1 linker), our LRET and tmFRET data identified position E35 and Y8 to be sitting in the window between the T1 domain and the transmembrane domain.

We propose an inactivation model for Shaker Kv channels where the inactivation peptide sits in the window between the channel and the T1 domain (Fig. 7). It interacts with the acidic residues of the T1-S1 linker. Fan and coworkers (13) proposed that the acidic residues stablilize the resting state position. During channel activation, the far end of the chain moves first, preceded by the movement of ball towards the pore. The final docking of the hydrophobic ball into the pore blocks the pore to bring the inactivation. Molecular dynamics of a Shaker IP have suggested that the IP adopts a helical structure which inserts spontaneously into the intracellular cavity of the pore and subsequently “snakes” deep into the pore when driven by voltage (51). It is possible that the slow fluorescence component of A3Anap and Y8Anap represent this last transition.

## Materials and Methods

### Xenopus oocyte injection

The mutations were introduced into the full-length Drosophila Shaker H4 gene in the pBSTA vector. The N-terminal stop codons (A3, Y8, K19, E35) were inserted into a 9LL background in which 9 noncanonical start codons have been removed via silent mutations as previously described (26, 27). For Anap and tco incorporation, 9.2 nl of 0.1 ng/nL cDNA encoding the tRNA/RS pair (22, 42) was injected into the nucleus of Xenopus laevis oocytes 6-24 hours prior to coinjection of 23 nL 1 mM Anap (Abzena TCRS, custom synthesis TCRS-170) or Tco (Sichem, SC-8008) and 35 ng in vitro transcribed RNA (23). All steps during and after Anap injection were performed under red light to avoid bleaching. Oocytes were then incubated for 1-3 days at 18°C in Barth’s solution supplemented with 5% horse serum. See Kalstrup & Blunck, 2017 (52) for detailed description and visualization of this procedure.

### Electrophysiology

Voltage clamp was performed with a CA-1B amplifier (Dagan). Currents were recorded in the cut open oocyte voltage-clamp configuration (53) and analyzed by using GPatch (Department of Anesthesiology, University of California, Los Angeles). For ionic recordings the external solution contained in mM: 5 KOH, 110 NMDG, 10 Hepes, and 2 Ca(OH)2, and the internal solution contained in mM: 115 KOH, 10 Hepes, and 2 EDTA. For gating recordings, the KOH was replaced by NMDG. All solutions were adjusted to pH 7.1 with MES. For 4-AP blocking experiments, 5mM NMDG (gating) or KOH (ionic) was replaced by 5 mM 4-AP, and the command potential was held at 0 mV for 5 seconds prior to recordings. Conductance (G) was calculated from the steady state currents (I) via G=I/(V-Vrev), where Vrev is the reversal potential. Conductance-, charge- and fluorescence-voltage relationships (GV, QV, FV) were fitted to a Boltzmann relation of the form y = (1 + exp (−(V − V1/2)/dV)) −1.

### Voltage clamp fluorometry

Fluorescence intensities were recorded with a photodiode detection system (Photomax 200, Dagan) using an ex-377/50, dc-409, em-470/40 filter set for Anap fluorescence. Bleaching effects during step protocols were accounted for by scaling fluorescence traces using the first sweep as reference.

### LRET measurements

For donor only labelling, oocytes were incubated in a depolarizing solution (115 mM K-MeSO3, 2 mM Ca(OH)2, 10 mM HEPES, at pH 7.1) containing Tb3+ complex (5µM) at room temperature for 20 min. For acceptor labelling, Cy3 tetrazine was injected into the oocytes a day before the experiment. The setup was based on a Zeiss Axiovert 200 microscope. Picosecond laser pulses of 337 nm from a pulsed 3-milliwatt N2-laser (Spectra-Physics) was directed in a wide-field illumination onto the oocytes containing the labeled channels. The light from the oocytes was collected using a 1.25 NA 40× glycerol immersion quartz objective (Partec) and filtered with a bandpass emission filter. Light was detected by a photon counter (Laser Components). Analysis of the lifetime decays was done with an exponential fitting program (MATLAB). Distances were determined using the following relations :

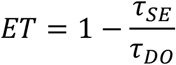

Where ET represents energy transfer, τ_DO_ is the time constant of the donor lifetime decay in absence of acceptor (donor-only), and τ_SE_ is the time constant decay of the sensitized emission of the acceptor.

The distance R between the donor and the acceptor was determined using:

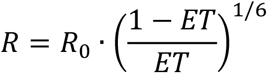

where R0 is the distance at which 50% energy transfer occurs. The R0 value used for the fluorophore pair of Tb3+ complex and Cy3 tetrazine was 61.2 Å (38).

### tmFRET measurements

Fluorometry experiments were performed on the VCF setup described above. CoSO4 was injected into the oocytes to a final concentration of 2µM for FRET experiment. Energy transfer efficiency was calculated according to the methods mentioned in Dai et al(45, 47).

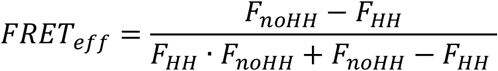

where FHH is the normalized fluorescence of channels with dihistidines and FnoHH is the normalized fluorescence of channels without dihistidines. The distances were then calculated using the Förster equation as follows :

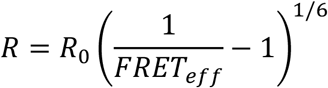

where R_0_ is the Förster distance for FRET between L-Anap and Co^2+^-dihistidine (12 Å).

## Data Availability

Data will be made available at a publicly accessible data repository.

## Acknowledgements

This work was financially supported by the Canadian Institutes of Health Research (CIHR, PJT-169160) and the Natural Sciences and Engineering Research Council of Canada (NSERC, RGPIN-2023-04752). CIRCA is a research center financially supported by the Fonds de recherche Québec — Santé.

## Supplementary Figure

**Fig S1.**
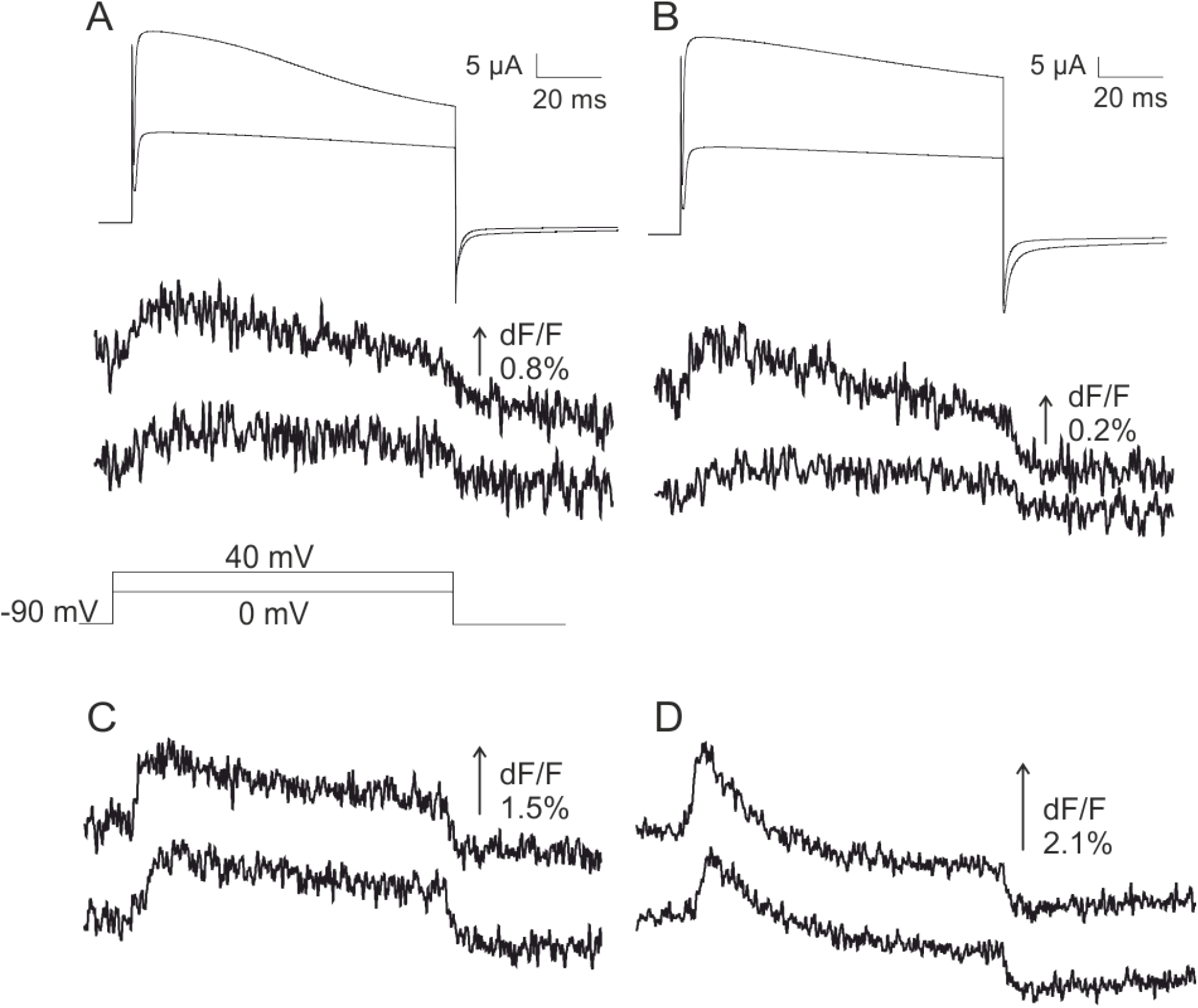
Two fluorescence components present in A3Anap and Y8Anap Currents and fluorescence changes elicited by depolarizations of A) A3Anap and B) Y8Anap channels, obtained at high expression levels. The same protocol was performed with C) A3Anap-W434F and D) Y8Anap-W434F channels

